# BECon: A tool for interpreting DNA methylation findings from blood in the context of brain

**DOI:** 10.1101/111609

**Authors:** Rachel Edgar, Meaghan J Jones, Michael J Meaney, Gustavo Turecki, Michael S Kobor

**Affiliations:** Department of Medical Genetics, University of British Columbia; Centre for Molecular Medicine and Therapeutics, BC Childrens Hospital; Ludmer Centre for Neuroinformatics and Mental Health, Douglas Mental Health University Institute; Singapore Institute for Clinical Sciences, Singapore; Canadian Institute for Advanced Research, Toronto, ON, Canada; Department of Psychiatry, McGill University

## Abstract

Tissue differences are one of the largest contributors to variability in the human DNA methylome. Despite the tissue specific nature of DNA methylation, the inaccessibility of human brain samples necessitates the frequent use of surrogate tissues such as blood, in studies of associations between DNA methylation and brain function and health. Results from studies of surrogate tissues in humans are difficult to interpret in this context, as the connection between blood-brain DNA methylation is tenuous and not well documented. Here we aimed to provide a resource to the community to aid interpretation of blood based DNA methylation results in the context of brain tissue. We used paired samples from 16 individuals from three brain regions and whole blood, run on the Illumina 450K Human Methylation Array to quantify the concordance of DNA methylation between tissues. From these data we have made available metrics on: the variability of CpGs in our blood and brain samples, the concordance of CpGs between blood and brain, and estimations of how strongly a CpG is affected by cell composition in both blood and brain through the web application BECon (Blood-Brain Epigenetic Concordance; https://redgar598.shinyapps.io/BECon/). We anticipate that BECon will enable biological interpretation of blood based human DNA methylation results, in the context of brain.

## Introduction

Research exploring the associations and underlying mechanisms of complex traits such as brain function and health have primarily focused on genetic variation, with some success^1–4^. Inter-individual variation in brain function and health emerges as a result of both genetic variation and environmental influences. Enduring effects of environmental exposures on brain function are of particular interest for our understanding of the origins of brain disorders and for the development of effective biomarkers. Epigenetic signals are an attractive candidate mediator of enduring environmental effects on cellular function. Indeed, there is now considerable evidence for the idea that environmentally-regulated epigenetic states might form the biological basis for gene x environment interactions^5–11^. DNA methylation (DNAm) is a relatively stable epigenetic mark that is amenable to genome-wide assessment in biosamples from human subjects in studies of complex traits. The model of DNAm as a mediator of complex traits has produced a surge of DNAm-based, epigenome-wide association studies (EWAS) in brain research with promising results. Studies of mammalian models of stress, anxiety, addiction and brain cell composition support the hypothesis that DNA methylation patterns are associated with brain function and health^12–17^. Moreover, there is evidence from human samples of specific patterns of DNAm linked to schizophrenia, autism, bipolar disorder and major psychosis^18–22^.

Tissue type is one of the strongest contributors to changes in methylation seen in EWAS ^23–25^, and therefore an important consideration in designing EWAS. Blood is a commonly used surrogate for brain in human studies due to accessibility and potential for direct relation to disease through hormonal and immune regulation^26–29^. However, due to the highly tissue specific nature of DNAm it is important to consider the concordance between tissues in order to interpret findings derived from surrogate tissues^23–25^. Indeed, previous work has demonstrated that genome-wide DNAm profiles are highly tissue-specific, both in terms of inter-individual variability and absolute measures^24,30–33^. In blood and brain specifically, using a Illumina 450K Human Methylation Array (450K) dataset of matched blood and brain tissues, we have shown that tissue identity, followed by cell type heterogeneity within a tissue, represent the largest contributors to DNAm variance^31^. Moreover, while human tissues share some common DNAm patterns associated with biological variables like aging^34^, tissues also show unique patterns in relation age^31^. Despite these findings, it remains largely unknown to what degree DNAm changes at individual CpGs found in blood can serve as biologically relevant indicators of human brain biology.

To enable the interpretation of DNAm results from blood based EWAS we aimed to quantify to what degree blood is informative of brain DNAm. Using genome wide analysis from paired human blood and brain run on the 450K we measured the level of concordance in DNAm across the methylome. Using variability and correlation thresholds on all CpGs, our results showed varying degrees of blood-brain concordance between CpGs. As we found concordance to be highly CpG dependent, blood based studies do indeed need to be carefully interpreted in the context of tissues specific DNAm differences. Therefore, as a tool for the community we have built a user friendly web application Blood-Brain Epigenetic Concordance (BECon) to explore blood based DNAm findings in the context of human brain

## Materials and Methods

### Data Collection

Using data from a previously published cohort, a total of 16 subjects were included in this study^31^ (one subject from the original cohort of 17 did not have a blood sample and therefore could not be used) (GSE).

### Quality Control and Normalization of 450K Data

Within Genome Studio samples were normalized by background subtraction and color correction, after which data was exported into R version 3.1.1. Sixty-five probes were removed as they directly measure SNPs and were not needed in this analysis beyond confirming replicate ID. Probes with evidence of cross-hybridization to regions other than the probes target in the genome were removed (41 937 probes)^35^. In addition, 1 035 probes were filtered as no calls (bead count less than 3) in 5% of samples and 1 342 probes were filtered as having 1% of samples with a detection p-value greater than 0.05. In total probe filtering left 441 198 probes in the processed 450K dataset.

Normalization was performed using BMIQ^36^, as quantro^37^ determined that quantile normalization would not be appropriate for this data set. The inappropriateness of quantile normalization was expected as our data set consisted of two very distinct tissues, with very different DNAm beta value distributions (Supplementary Figure S1).

The 63 samples (4 samples from 15 subjects and 3 samples from the one subject missing BA20), were examined with principal component analysis (PCA) to visualize the presence of batch effects. In this study the first principal component (PC) is not shown in visualizations (see Supplementary methods). The loadings of each PC were associated with technical and biological variables using ANOVA for categorical variables or Spearman correlations for continuous variables. Array barcode as well as refrigeration delay and sample pH showed strong associations with the top PCs loadings on samples (Supplementary Figure S2). ComBat was used to remove the batch effects of array barcode, refrigeration delay and PH from the DNAm data^38^.

### Cell Composition

As the dataset was comprised of blood and brain samples, cell composition was estimated for each of the two tissue types. Specifically, blood cell type proportions were estimated based on reference epigenomic profiles for six major sorted cell types^39^ and normalized between blood samples^40^. Similarly, neuron and non-neuronal brain cell type proportions were estimated^41^ and normalized between brain samples^40^(Supplementary Figure S3).

As a check of the pre-processing steps, root mean squared errors (RMSE) were calculated between replicates. Root mean squared errors between replicate pairs remained high across all stages of quality control and pre-processing, indicating normalization and batch correction successfully removed noise from the data (Supplementary Figure S4).

### Differential DNA Methylation Analysis

Mean DNAm across samples of a tissue were correlated between all tissue pairs using Spearman correlations as a DNA methylome-wide indicator of general sample similarity. To quantify the similarity of tissues at individual CpGs, differential DNAm analysis at each CpG was performed between all tissues pairs. Linear models were run with covariates for subject gender and age, and to account for the paired structure of the samples, subject ID was included in the model. Multiple test correction was done on the nominal p values of each tissue pair comparison, using Benjamini-Hochberg correction ^42^.

### Informative CpG selection

The correlation of DNAm level at each CpG was calculated between blood and brain separately for each brain region brain using Spearman correlations on M values. The variability of a CpG across individuals was measured as the range between the 10th percentile and the 90th percentile of blood sample CpG betas^43^. This reference range is intended to capture variability in the bulk of the samples while limiting the effect of outlier measures at a CpG, which would otherwise give a falsely high estimate of variability.

To define informative CpGs we used biologically relevant thresholds for both correlation and variability. The most highly correlated and variable CpGs observed were the polymorphic CpGs (CpGs with a known SNP at the cytosine or guanine of the CpG) and those on the X and Y chromosomes (presumably highly variable because the cohort contains males and females). While these 32 344 CpGs were not of explicit interest in this study, and later removed from analysis, the variability and correlation distributions of these 32 344 highly correlated polymorphic and sex chromosome CpGs were used to guide the selection of the correlation and variability thresholds for informative CpGs. A full explanation of the threshold selection is provided in the supplementary text, but in short, informative CpGs had to meet a variability threshold of 0.1 reference range and correlation threshold of 2 standard deviations from the mean correlation of the highly positively correlated polymorphic and sex chromosome CpGs enriched correlation peak, defined separately for each brain region.

The importance of a variability threshold was clear when the matched blood and brain samples were randomly unmatched in five simulations. The correlation distributions of these unmatched data sets were used as null distributions to compare to the real paired data correlation distributions ^44^.

### CpG to Gene Associations

There are multiple approaches for associating a CpG to a gene, such as the closest TSS^35^, associating CpGs to gene by the CpGs localization to a genes body or promoter^45^, or stringent associations based on CpGs with only one likely gene association (i.e. lone gene associations)^31^. These approaches focus on a single gene, rather than allowing for multiple gene associations for a CpG. We used a CpG to gene association definition that allows for a CpG to be associated with multiple gene features, as well as multiple genes (see Supplementary text). This inclusive association, while somewhat more ambiguous, is an attempt to capture all possible roles of a CpG in gene regulation^46^. The gene list used was the Refseq genes from UCSC, including all splice variants of Refseq mRNA. The gene list included 24 047 genes and a total of 33 431 unique transcription units. The 485 512 CpGs on the 450K array associated with 23 018 genes (43.8% intragenic CpGs, 34.2% promoter CpGs, 2.5% 3 region CpGs, and 19.5% intergenic CpGs)^46^.

### Comparison to Previous Analyses

DNAm in blood and brain samples was analyzed previously using a 450K dataset^33^ providing an opportunity to validate our findings using an independent dataset. The published data set provided 74 matched brain and blood samples on Gene Expression Omnibus (GEO; GSE59 6 85)^33^. The regions examined in this previous work were cerebellum, entorhinal cortex, frontal cortex and superior temporal gyrus regions. While their frontal cortex and our BA10 region partially overlap, the 3 other regions available in GSE59685 allow for possible validation in structurally and functionally different brain regions. We ran the GSE59685 normalized data through our pipeline (described above) to make the results as comparable as possible. Unfortunately, in GSE59685, the tissues were run on separate arrays, introducing confounding of array and tissue. However, despite this limitation, to be consistent with our analysis we did run ComBat to correct for sentrix ID. Cell correction was performed as described above in the brain regions and blood. Spearman correlations and reference ranges were calculated between blood and all brain regions, and informative CpGs were defined similarly as described above. The actual percent overlap of informative CpGs was calculated for all 7 brain regions available. Monte Carlo simulations were used to build an expectation of overlap between two lists of informative CpGs.

To test for enrichment of mQTL in informative CpG lists we used mQTL previously identified at p¡11014 in middle aged individuals using the mQTL database^47^ (http://www.mqtldb.org/). The mQTL list contained 31 325 CpGs under observed genetic influence. Using Monte Carlo simulations we built an expected overlap of the mQTL CpGs and informative CpGs, to compare to the observed overlap.

There have been numerous studies using blood as a surrogate for brain when studying a neurobiological disorder^48^. To explore whether CpGs identified in these studies are informative of brain DNAm we collected a list of six CpGs associated with 4 genes previously observed to be differentially methylated in blood in relation to psychiatric disorders^48^. We explored the identified CpGs correlation between blood and brain in BECon. Specifically, the CpGs were evaluated in terms of the correlation percentile in each brain region to explore if the CpGs are more or less informative then the average CpG. CpGs we also examined for variability in each tissue as a measure of biological relevance.

## Results

### Differential Methylation Analysis of Blood and Brain

Tissue is the one of the largest contributor to DNAm variance^31,49^. However, it is not yet known which specific CpGs show concordance in DNAm between blood and brain, at which blood DNAm could potentially serve as a proxy for brain DNAm, and which CpGs show no concordance. DNAm was measured from matched human blood and brain on the 450K^45^. In comparisons of DNAm between the four samples from each of the 16 individuals across the methylome, different brain regions from the same individual had higher correlation with each other than any brain region with blood (Figure 1A). Additionally, brain to blood DNAm analysis returned orders of magnitudes more differentially methylated CpGs than did differential DNAm analysis between brain regions (e.g. 119 371 differential CpGs between BA10:blood and 347 between BA10:BA7, FDR¡0.001, mean difference in DNA methylation between tissues 0.1; Figure 1B).

**Figure 1:**
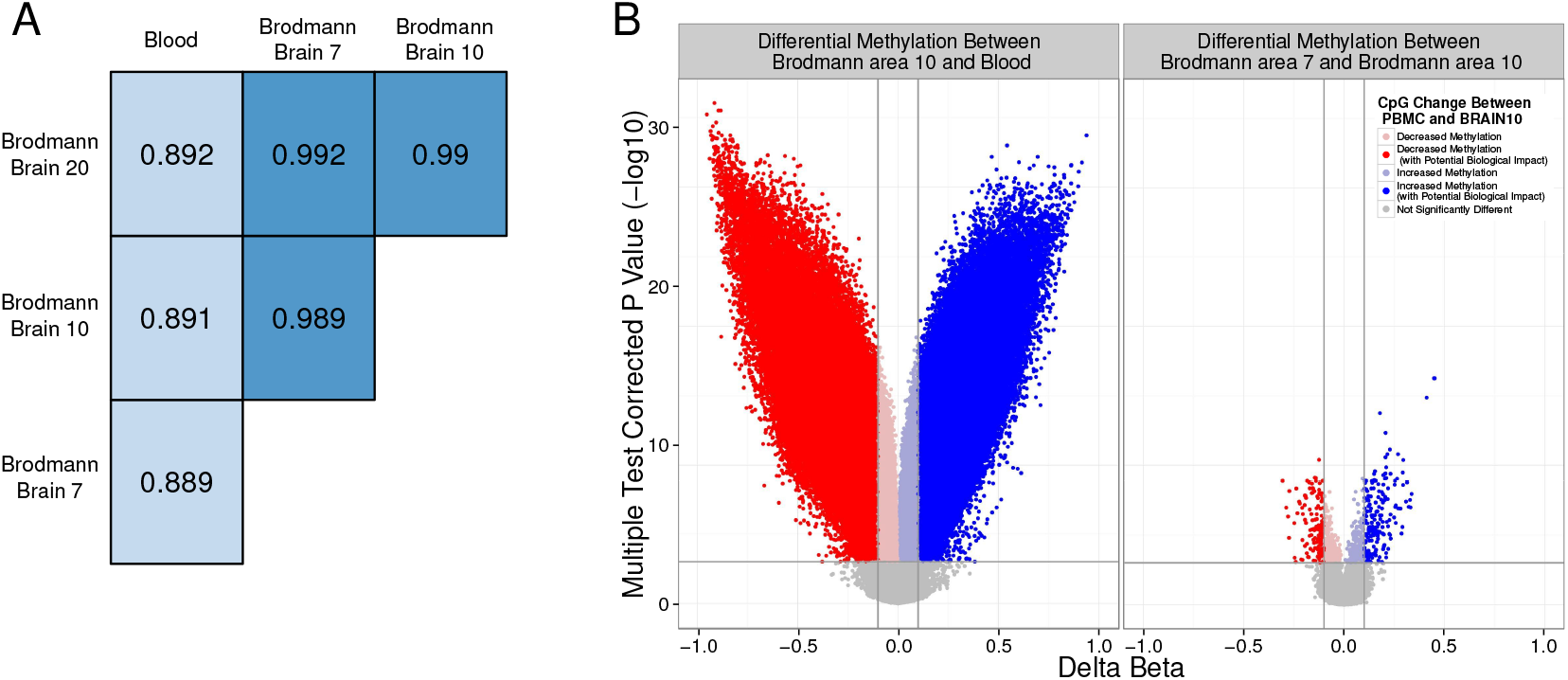
Human blood and brain show very distinct methylation patterns. A) DNAm correlation values between each tissue pair from an individual, averaged across all individuals. B) Volcano plots of the differential methylation analysis between representative tissue pairs (blood and Brodmann area 10; and Brodmann area 10 and 7). Vertical lines indicate a DNAm difference between compared tissues of 0.1. The horizontal line represents an FDR corrected p value of 0.001. Points are coloured to highlight CpGs exceeding both the biological and statistical cutoffs.

### Informative CpGs Exist between Human Blood and Brain

While our group and others have observed large differences between human blood and brain DNAm^23,31^, by necessity, blood is often used as a surrogate in for brain tissue. We therefore set out to use the strength of our matched sample cohort to identify CpGs that show concordance between blood and brain. Our first step in identifying these CpGs was to use the correlation of inter-individual variability between blood and each brain region (Figure 2A). These correlations had a slightly skewed distribution toward negative correlations, indicating the majority of CpGs are not concordant between blood and brain, but a few CpGs are highly positively correlated (Supplementary Figure S5; see Supplementary Text).

**Figure 2:**
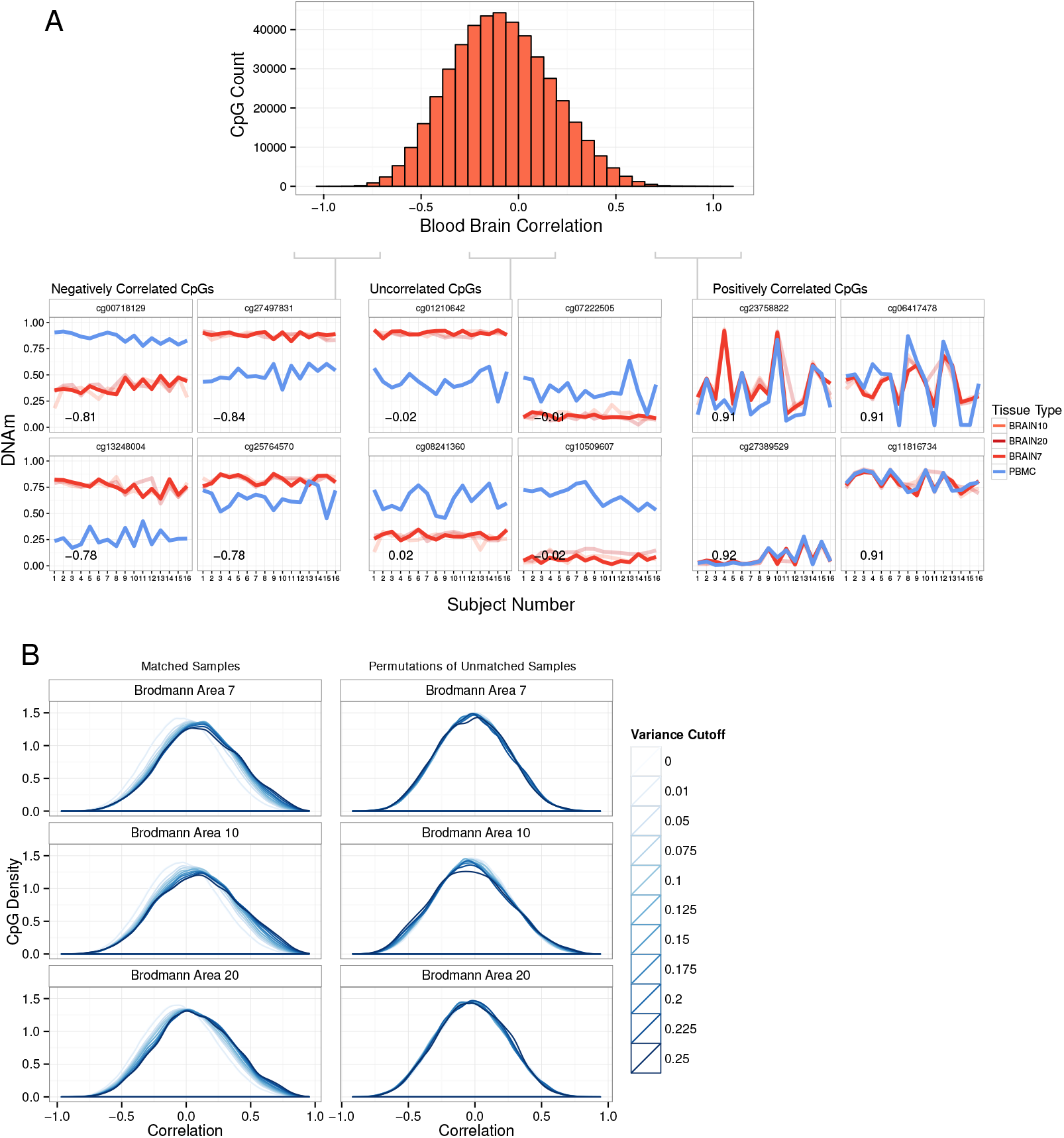
Paired tissues allow the identification of blood-brain concordance across individuals. A) Representative distribution of all CpG's correlation values between blood and Brodmann area 7. Line plots show DNAm of representative CpGs from three sections of the correlation distribution. The line plots show the correlation of inter-individual variability at the representative CpGs. Lines are colored by tissue and BA10 and BA20 are shown fainter. B) Correlation distributions of CpGs are shown for each brain region at increasingly stringent variability cutoffs. Line colours darken as the CpGs underlying the distribution become more strictly thresholded on reference range. Plots on the left show the distributions of matched blood and brain correlation values, and on the right the distributions are for unmatched permutations of sample order to simulate unpaired data.

However, many of the most highly correlated CpGs had very low variability between individuals, which likely limits the utility of these CpGs to explain differences in phenotype and/or exposures in EWAS. We therefore explored the importance of variability in defining concordant CpGs. We looked at the correlation distributions of increasingly more variable sets of CpGs (Figure 2B) and found that the higher the variability threshold the more skewed to positive correlations the distribution became. This trend was not an artifact of the variability measurements as it disappeared when the data was simulated as unpaired (Figure 2B). We therefore endeavored to select CpGs with both high inter-individual variability, as well as high blood-brain correlation.

To make the thresholds of variability and correlation more biologically driven we based the thresholds on a set of CpGs which were some of the most highly variable and correlated between blood and brain, polymorphic CpGs and CpGs on sex chromosomes (Figure 3A). We focused on variability in blood and blood-brain correlation of these 32 344 CpGs to define our thresholds. The reference range variability of these CpGs in blood was 0.11 so a threshold of 0.1 was used for our concordant CpG selection (Figure 3B). The correlation distribution of all CpGs was bimodial, and we focused on the polymorphic and sex chromosome enriched highly positively correlated peak to define our correlation thresholds (Figure 3C). Therefore our definition of a blood-brain informative CpG is a CpG at which the DNAm in blood correlated with DNAm in brain (rs = BA7 0.36; BA10 0.40; BA20 0.33) and DNAm is also highly variable in blood (reference range ¿0.1).

**Figure 3:**
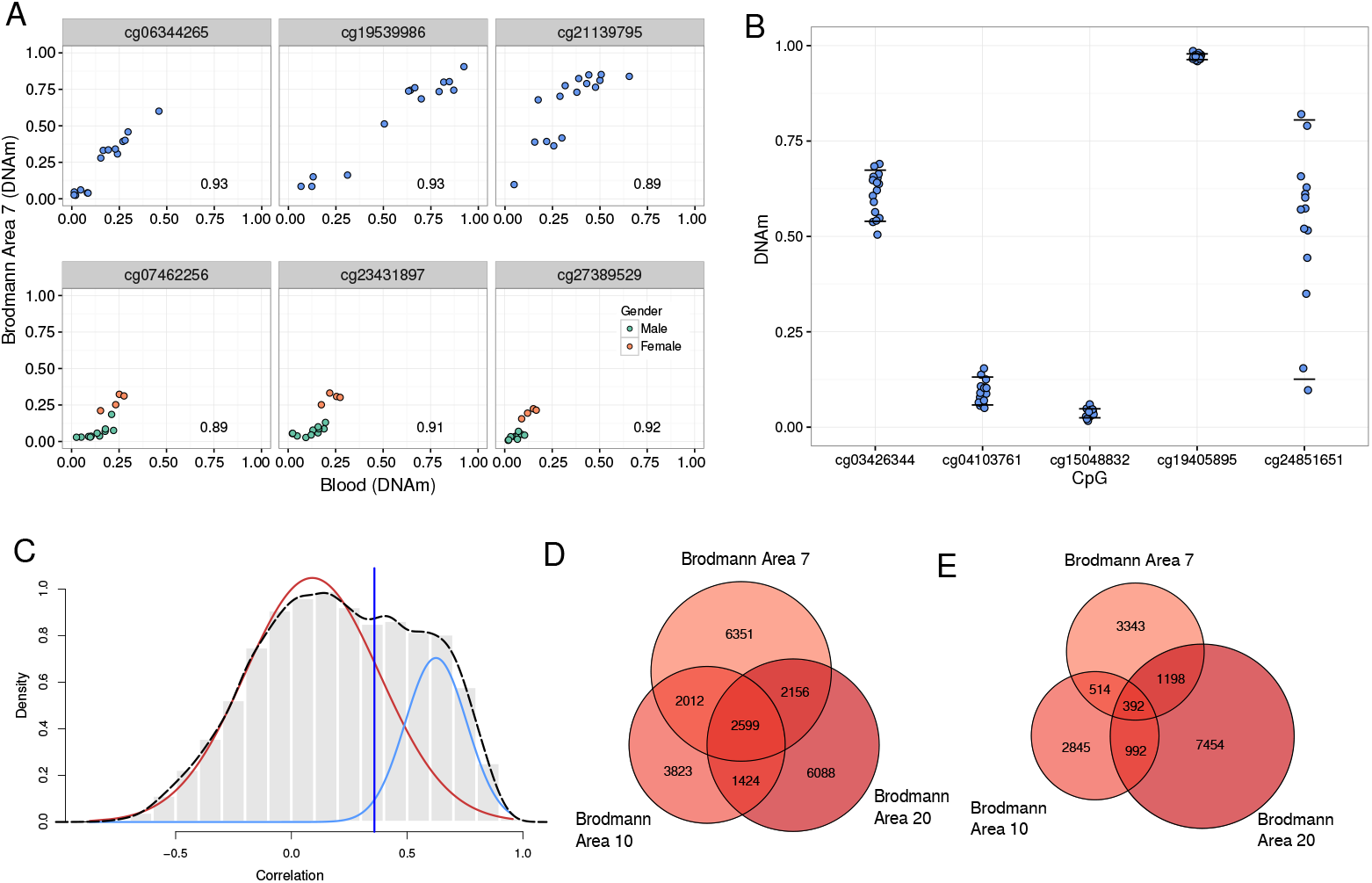
Informative CpGs are defined through both variability and correlation thresholds. A) Representative SNP and sex chromosome CpGs show high levels of correlation and variability. All plots show the relationship between DNAm in Brodmann brain area 7 and blood with the correlation coefficient in the bottom right of the plot. The CpGs in the top 3 plots show a polymorphic CpG. The CpGs in the bottom three plots show sex chromosome CpGs, and individuals are coloured by gender. B) Reference range variability in blood DNAm is shown at representative CpGs. Horizontal lines at each CpG represent the 90th and 10th percentile of blood DNAm level. Reference range is the range between the horizontal lines. C) Definition of the correlation coefficient threshold for informative CpGs in Brodmann brain area 7. The histogram and broken line show the correlation distribution for CpGs passing the strictest variability threshold. Solid lines are the 2 fitted Gaussian components of the distribution(red generally uncorrelated peak, blue positively correlated peak). The vertical black line indicates 2sd away from the positively correlated peak mean which was used as the correlation threshold for blood-brain informative CpGs. D) Venn diagram showing the overlap of informative CpGs with positive correlations between blood and brain in the three Brodmann areas sampled. E) Venn diagram showing the overlap of informative CpGs with negative correlations between blood and brain in the three Brodmann areas sampled.

Using our variability threshold of 0.1, we identified 83 427 variable CpGs. Of these, 48% also passed our correlation requirements, resulting in a total of 40 029 informative CpGs (both positively and negatively correlated). Thus 9.7% of the total number of CpGs examined were informative between blood and any of the three brain regions. Informative CpGs identified in each brain region show a large overlap (Figure 3D and E), as expected considering the observed similarities in DNAm of the three brain regions. While we have been strict in our definition of what an informative CpG is, and small changes to the correlation and variability thresholds do result in large changes for the number of CpGs considered informative (Supplementary Table S1), our informative list has been built on biologically defined statistical thresholds based on the polymorphic and sex chromosome CpGs characteristics. Therefore our informative CpG list is relatively stringent and should reflect the strongest concordance signal in the data.

### Informative CpGs were Enriched in Intergenic Regions of the Genome

Next, we explored the genomic location(*s*) of our informative CpGs. We associated each CpG with a gene (see Supplementary Text), and used the Illumina annotation for CpG island associations ^45^. In general, informative CpGs were depleted in gene promoters and CpG islands and enriched in intergenic regions (p¡0.001; Figure 4).

**Figure 4:**
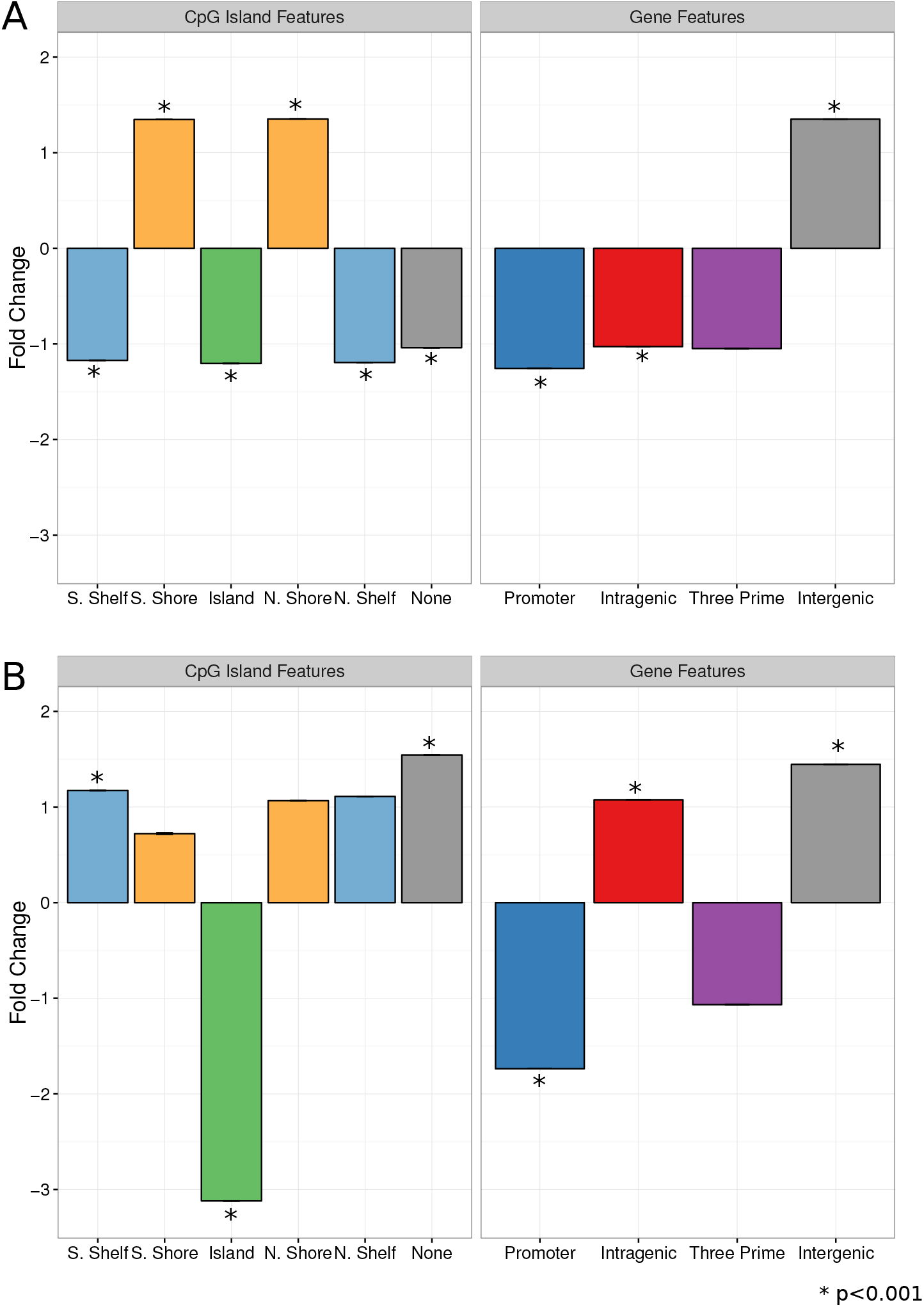
Blood brain informative CpGs show associations to specific genomic features. In all plots, bars show the fold change between informative CpG count in each region and the count of CpGs from 10,000 permutations of random CpGs in that same region. Error bars show standard error. A) Genomic enrichment for informative CpGs which are positively correlated between blood and brain. B) Genomic enrichment for informative CpGs which are negatively correlated between blood and any brain.

We then explored whether genes associated with informative CpGs were involved in any specific biological process. We used a list of 239 genes associated with at least ten informative CpGs (informative genes), in order to focus on the genes with high DNAm variability and concordance between blood and brain. The informative gene list showed enrichment for GO terms related to cell-adhesion and highly multifunctional genes (Supplementary Table S2)^50^. We then investigated if the informative genes were more highly expressed in either blood or brain, using independent gene expression datasets (GSE17612, GSE37171 and GSE61635). The informative genes are less expressed in blood samples than the average expression of all genes measured (p¡0.001, Wilcoxon Rank Sum) but the expression of the informative genes was not different from the average expression of all genes in brain samples (p=0.98, Wilcoxon Rank Sum; see Supplementary Text and Figure S6). Therefore, informative genes may be more brain specific than blood specific as they are less expressed in blood then the average gene.

### Comparison of Informative CpGs to Previous Findings

An existing similar blood brain DNAm analysis^33^ provided an opportunity for independent validation of our results. The previous study, reports the correlation between blood and four brain regions (cerebellum, entorhinal cortex, frontal cortex and superior temporal gyrus), and provides data for 74 individuals with paired blood samples DNAm (GSE59685). We found that while the greatest overlap of informative CpGs was between brain regions from the same study, the overlap between the lists of informative CpGs from the two studies was greater than expected by chance (Figure 5A). Interestingly, the previous study had very few negatively correlated sites (42 negatively correlated informative CpGs compared to 16 738 in our data set), which may suggest the negative correlations we observed represent a property inherent to our smaller sample size (Supplementary Figure S7).

**Figure 5:**
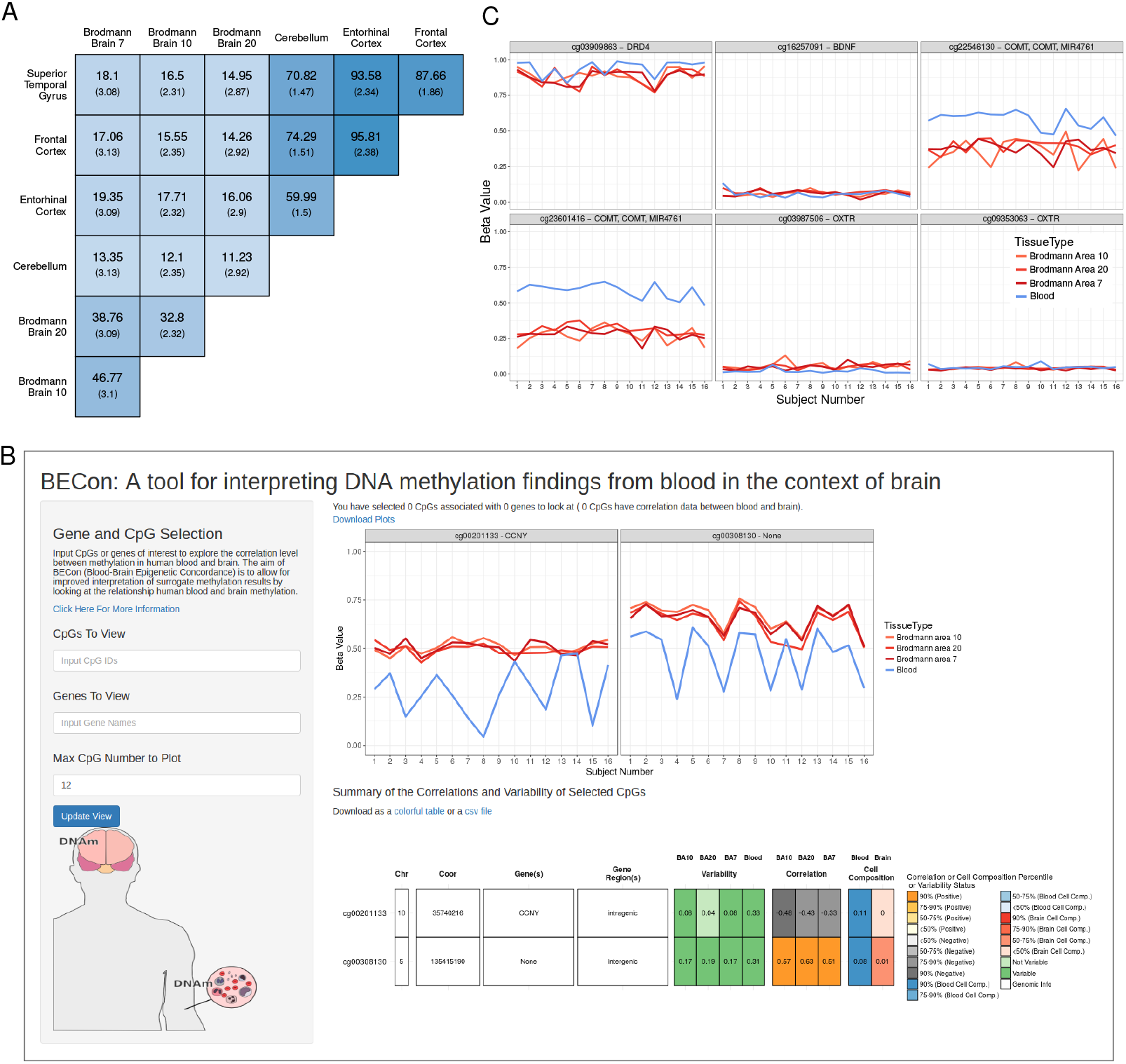
The informative CpG list can be used to validate previous findings. A) Percent of overlapping positively correlated informative CpGs in each of our brain regions and those regions used in the Hannon et al. (2015) data. The number in larger text is the percent of the informative CpGs along the rows that are also informative in the tissue along the columns, the smaller number is the percent overlap of these lists expected by chance. The color of each box is based on the percent overlap. B)Visualization provided in BECon. The plots show the inter-individual variability at two representative CpGs. The table shows the various metrics provided in BECon for each CpGs queried. C) Inter-individual variability at the CpGs identified previously in blood based studies of psychiatric disorders.

As polymorphic CpGs were some of the most highly concordant CpGs in our data, we speculated that our informative CpGs could be under genetic influence and represent potential DNAm quantitative trait loci (mQTL). We tested whether our informative CpGs and those we defined in GSE59685 were enriched for known mQTL using the mQTL database^47^ (http://www.mqtldb.org/). We found 8 202 out of 40 029 (21%) of our informative CpGs and 3 018 out of 10 930 (28%) informative CpGs from the Hannon et al. (2015) cohort were previously identified as mQTL. In both informative lists the mQTL numbers represented significant enrichment for known mQTL (Monte Carlo simulations, pj 0.0001). Although our smaller cohort likely did not have enough genetic diversity to capture all potential mQTL sites (Supplementary Figure S8), our informative CpGs were still enriched for mQTL.

### BECon as a Community Resource

The availability of this matched blood and brain DNAm data set provided an excellent opportunity to develop a community resource. We have built an R Shiny web application^51^ to aid interpretation of blood based DNAm results in studies of brain function and health (Blood-brain Epigenetic Condordance; BECon; https://redgar598.shinyapps.io/BECon/). While we also made the full data set available on GEO (GSEnumber), we built BECon for researchers interested in a particular gene or CpG, but without the need to re-analyze our data themselves. A full description of the information provided through BECon is in the supplementary text, with detailed explanations for how the metrics provided were calculated. To aid interpretation of epigenetic results BECon provides metrics on: the variability of CpGs in our blood and brain samples, concordance DNAm at CpGs between blood and brain, and estimations of how strongly a CpG is affected by cell composition in both blood and brain (Figure 5B).

To demonstrate the utility of BECon, we assessed key candidate genes often investigated for changes in DNAm in relation the psychiatric disorder (BDNF, COMT, OXTR and DRD4). We examined six specific CpGs in these candidate genes that were identified as differentially methylated in blood in studies of psychiatric disorders^48^ (Supplementary Table S3). The six CpGs show varying levels of average correlation across brain regions (rs=-0.15 − 0.49) and one CpG is in the 90th percentile of all CpG correlation values (Figure 5C; Supplementary Table S4), suggesting only some differential DNAm reported previously in blood could be expected to be seen in brain.

## Discussion

The increasing popularity of EWAS using blood samples to study brain function and health outcomes has created a need for tools to enable interpretation of DNAm results in the context of the brain. Our findings indicate that it is essential to examine the concordance of DNAm between blood and brain at each CpG before interpreting blood-based results, as concordance varied greatly dependent on CpG. A subset of CpGs, which we consider informative of brain, has been validated in an independent cohort. Despite the tissue specific DNAm seen between blood and brain the validated informative CpGs suggested blood has applicability as a surrogate for brain.. In identifying informative CpGs, correlation seemed to be the most logical measure of concordance, however we found a variability measure was also necessary to identify concordance with the utility to explain differences in phenotype and/or exposures in EWAS. Discordant CpGs were either not variable in one tissue, or appear to be potentially tissue specific in their variability. Tools to examine multiple metrics of concordance simultaneously will aid the interpretation of blood based DNAm results. We developed BECon to enable easier examination of the concordance of blood and brain DNAm. Our hope is that BECon that should allow for biologically grounded interpretation of blood-based DNAm results.

The web application BECon that we provide to the community includes metrics on the variability of CpGs in blood and brain. We have included metrics on variability as we found that the DNAm at many correlated CpGs varied by less that would generally be considered biologically meaningful. It is the consensus of the field that differences in DNAm, whether between tissues or disease states, to be have a theoretical biological impact they must show variability in DNAm measures^52^. When examining concordance of DNAm in BECon it is expected that if a CpG shows no variability in either blood or brain tissue, that any concordance will not be biologically meaningful. While CpGs not variable in our blood and brain data set may vary in another context, or perhaps another tissue, we are hopeful that or concordance findings are robust and relevant to other blood and brain samples.

We have some evidence our concordance findings are robust to brain region type and cohort as we were able to confirm the general trends seen in previous blood-brain studies^33,53^. Our study, like others, demonstrated that tissue type is a considerable contributor to DNAm variability, evident by the abundant DNAm differences we have observed between each brain region and blood. Second to broad tissue type differences the next largest contributor to DNAm variation is cell composition within a tissue^31^. In studies using surrogate tissues, as with all DNAm studies, it has become more apparent that cell type composition adjustments are a mandatory step in the analysis of data for results to be interpretable. We therefore included statistics on the estimated effect of cell type composition in BECon to better enable researchers to examine the effect of cell composition on CpGs or gene of interest.

Next to tissue type and cell composition, genetics are a major contributor to DNAm variability. CpGs under the influence of SNPs are some of the most variable CpGs observed in DNAm^47,54,55^. Given our variability selection criteria it is not surprising that out informative CpGs were enriched for mQTL. However it is reasonable to suspect that, regardless of variability criteria, CpGs under genetic control will be the most concordant between blood and brain as the genetics will be consistent between tissues. When interpreting DNAm concordance between blood and brain through BECon it will be important to consider the biology driving the concordance. At some CpGs the driver of variability and concordance may be primarily genetic and potentially independent of any environmental influences seen in associated with blood DNAm. While we were able to observe enrichment for mQTL we speculate that due to our smaller sample size we were not as sensitive to detect mQTL as the previous blood brain analysis^33^. With 16 individuals we may not have had representative samples from all possible alleles at many potential mQTL SNP loci. Therefore DNAm at potential mQTL was less variable in our data and high correlations could not be observed. Interestingly, this suggests that there is a minimum required sample size for thorough mQTL detection above 16 individuals, however we can not speculate if the 74 individuals used previously were enough to detect all possible mQTL.

Future studies into the gene regulation and expression associations of concordant CpGs in blood and brain may provide insight into the biological relevance of blood DNAm in brain. While we were unable to directly compare the concordance of our sites in gene expression data we were able to look at the overall expression of our informative genes in brain and blood. Interestingly we found that informative genes are less expressed in blood than expected by chance. It is possible that by our selection criteria we have identified CpGs that can serve as biomarkers for brain specific genes that have little to no function in blood and are therefore not highly expressed.

In addition to the highly tissue specific nature of DNAm, using a surrogate tissue for brain DNAm is further complicated by the existence of higher levels of hydroxymethylation in the brain compared to other tissues^56–58^. Hydroxymethylation has been seen in the brain at levels as high as 0.65% but only at 0.027% in blood^57,59^. Here we have characterized the anticipated utility of blood as a surrogate for brain in terms of a composite DNAm signal (mC+hmC). Hydroxymethylation in the human brain has potentially added complexity to our data and to the correlations calculated between blood and brain.

Using BECon we were able to identify a CpG showing high concordance between blood and brain that has also been identified as differentially methylated in a study of blood from patients with schizophrenia and controls (cg03909863)^29^. The CpG is a promising candidate to show DNAm concordance in the brains of the individuals, and could be relevant functionally in the brain as the CpG in located in coding region of dopamine receptor D4 (DRD4). In future studies that use blood as a surrogate for brain, BECon will enable prioritization of CpGs for validation to those CpGs with demonstrated concordance between blood and brain.

Despite the limitations of our analysis, there is a tremendous value and information content in quantifying the genome wide concordance of DNAm in the blood and brain. In anticipation of the communitys interest in examining whether specific CpGs in blood are informative of brain DNAm, we have built BECon to enable examination of concordance between blood and brain. We expect BECon to be most useful to users who need to interpret blood DNAm results from a study of brain function and health. This application may also help guide blood based surrogate studies toward candidate gene approaches or post-hoc selection of CpGs for validation.

## Acknowledgements

We would like to thank Dr. Magda Price, Sarah Goodman, Sumaiya Islam and Jack Hickmott for comments on the analysis and manuscript. This work was supported by: R. Howard Webster Foundation (F13-00031 to MSK); W. Garfield Weston Foundation/Brain Canada Foundation (F13-02369 to MSK MJM and GT). MSK is a Canada Research Chair in Social Epigenetics and the BC Leadership Chair in Child Development.

## Conflict of Interest

The authors declare no conflict of interest.

